# A novel dimerization site in non-structural protein 5A of hepatitis C virus regulates viral replication fitness

**DOI:** 10.64898/2026.05.05.722906

**Authors:** Paul Rothhaar, Thibault Tubiana, Colin Förster, Gabriel Vanegas Arias, Tomke Arand, Noemi Schäfer, Philipp Ralfs, Christian Heuss, Antonio Piras, Andreas Pichlmair, Xavier Hanoulle, Stéphane Bressanelli, Volker Lohmann

## Abstract

We previously found that high genome replication fitness of the hepatitis C virus (HCV) was associated with severe disease in immunocompromised patients. Elevated replication fitness was mediated by accumulation of mutations in the replication enhancing domain (ReED) within domain (D) 2 of non-structural protein (NS) 5A. NS5A is a partially unstructured phosphoprotein lacking enzymatic activity but fulfilling a key role in HCV replication due to interacting with various cellular and viral proteins. It can exist in a variety of dimeric and oligomeric conformations mediated by NS5A D1 with clinically approved NS5A inhibitors proposed to exert their antiviral function by fixing these dimers in distinct conformations. In this study, we aimed at elucidating the ReED’s mode of action. AlphaFold modelling indicated a so far unrecognized NS5A dimerization site in the ReED. Indeed, split nano luciferase assays revealed a significantly stronger NS5A dimerization of high replicator ReED variants, suggesting that high replication fitness is mediated by enforcement of NS5A self-interaction. This hypothesis was supported by the effect of low dose (1 pM) NS5A inhibitor treatment, increasing replication fitness and phenocopying the effects of ReED mutations. Furthermore, we found that HCV isolate JFH1, replicating with very high efficiency, is completely resistant to the regulatory function of the ReED. Chimeric replicons composed of ReED resistant JFH1 and the ReED sensitive isolate J6 identified NS3 helicase and NS5B polymerase as critical genetic elements mediating ReED sensitivity/resistance.

Our data overall suggest that NS5A is a negative regulator of HCV replication fitness with dimerization releasing the inhibitory interaction with helicase and/or polymerase, thereby likely facilitating initiation of RNA synthesis.

## Introduction

The hepatitis C virus (HCV) causes chronic infections in 60-80% of infected patients (1). Modern direct acting antivirals (DAAs) have cure rates >95% (reviewed in (2)) but lack of both access to treatment and a prophylactic vaccine lead to HCV still being a public health issue with over 50 million infected in 2022 (3). HCV has high genetic heterogeneity and is therefore classified into 8 genotypes (gts) with over 100 subtypes (4). The HCV genome has a length of 9.6 kb and is a positive sense single stranded RNA with one open reading frame (ORF) flanked by untranslated regions (UTRs). The resulting polyprotein is cleaved into 10 viral proteins by host and viral proteases. The 3 structural proteins core, E1 and E2 form the viral particle. Of the 7 non-structural (NS) proteins p7 and NS2 are crucial for particle formation but dispensable for genome replication which is performed by the other 5 NS proteins NS3-NS5B together with the UTRs (reviewed in (5)). NS3 functions both as a helicase and together with the co-factor NS4A as a protease. NS4B is needed for membrane rearrangements to form the HCV replication organelle. NS5A is a phosphoprotein lacking enzymatic function interacting with a variety of host and viral proteins. NS5B is the RNA dependent RNA polymerase (RdRp).

Having a defined viral replicase is key to study HCV replication fitness in cell culture with the help of subgenomic replicons (SGRs). Here, the replicase is expressed together with a luciferase reporter allowing for the detection of HCV replication (reviewed in (6)). Initially, replicating HCV wildtype (WT) isolates in the hepatoma cell lines commonly used in cell culture failed, with the exception of the gt2a isolate JFH1 (7). For all other isolates, only the virus acquiring cell culture adaptive mutations enabled replication (8, 9), eventually allowing for DAAs being developed based on the replicon system. More recent studies revealed that the presence of the cytosolic lipid transporter SEC14L2 (10) or pharmacological inhibition of the host factor Phosphatidylinositol 4-kinase III alpha (PI4KA) enables HCV WT replication in cultured hepatoma cells (11). The latter mimics cell culture adaptive mutations found in NS5B and NS5A (11). The crucial role of NS5A is further underlined by the fact that beyond viral protease and RdRp inhibitors, the third class of clinically approved DAAs are NS5A inhibitors (2).

NS5A consists of an amphipathic helix anchoring it to membranes and three domains (D) separated by low complexity sequences (LCS) (12). D1 is structured and allows for NS5A dimerization and potentially oligomerization which are crucial for NS5A functionality (13–15). The remainder of the protein is largely unstructured (16, 17). D1 and D2 are required for the RNA binding capability of NS5A (18) while the main phosphorylation sites can be found in LCS1D2 and D3 (reviewed in (19)). D3 is also crucial for viral particle formation (20). In LCS1D2, we previously characterised the replication enhancing domain (ReED) as a regulator of genome replication fitness (21). The norm appears to be a ReED allowing for low levels of replication with mutations leading to higher replication fitness mainly emerging in immunocompromised patients leading to severe disease in liver transplant patients (21). The sequence signature of elevated replication fitness was 3 or more mutations in the interferon sensitivity determining region (ISDR) (21) where mutations were also associated with response to antiviral interferon therapy (22).

While we could previously highlight the strong, pangenotypic impact of mutations in the ReED on genome replication fitness (21), it remains unclear how these mutations actually affect NS5A function. In the following, we aimed at clarifying the ReED’s mode of action in HCV replication enhancement. We could show that the presence of different ReEDs did not alter the interaction with host factors. However, AlphaFold modelling revealed a so far overlooked dimerization site in NS5A D2 which was confirmed by in vitro dimerization assays, with high replicator ReEDs leading to stronger NS5A dimerization. The ReED’s function was affected by determinants in NS3 and NS5B which overall suggests that the ReED acts directly of the HCV replicase.

## Results

### The ReED does not significantly affect NS5A interactions with cellular proteins

We recently described that accumulation of mutations in the ReED within NS5A could drastically enhance viral genome replication fitness (21). Nevertheless, it remained unclear how ReED mutations acted mechanistically. NS5A lacks enzymatic activity and exerts its functions in the HCV life cycle through interacting with viral and cellular proteins with interaction sites with protein kinase R (PKR) (23) and cyclophilin A (CypA) (24) being localised in the ReED (Fig. 1A, upper panel). To assess a connection between replication enhancing mutations in the ReED and alterations in host factor interactions, we performed an NS5A immunoprecipitation (IP) followed by mass spectrometry. We focussed on gt1b prototype isolates: the low replicator Con1 (25), the high replicator GLT1 (26) and the highly replicating Con1 based chimera harbouring the GLT1 ReED (21). To maximise IP efficiency and to generate comparable data, we inserted an HA-tag into NS5A at a well characterized insertion site in D3 (27) and performed immunoprecipitation using anti HA-beads. Using subgenomic replicons containing the viral replicase (NS3-5B), the UTRs and a luciferase reporter (Fig. 1A, lower panel) transfected into the hepatoma cell line Huh7-Lunet overexpressing the lipid transporter SEC14L2 (10), we could show that the HA tag had no major impact on replication (Fig. 1B). IP experiments were performed using constructs containing the T7 promotor leading to continuous protein expression in Huh7-Lunet SEC14L2 cells expressing T7 polymerase, maximising protein yield independent of replication fitness (Fig. 1C).

**Fig. 1:**
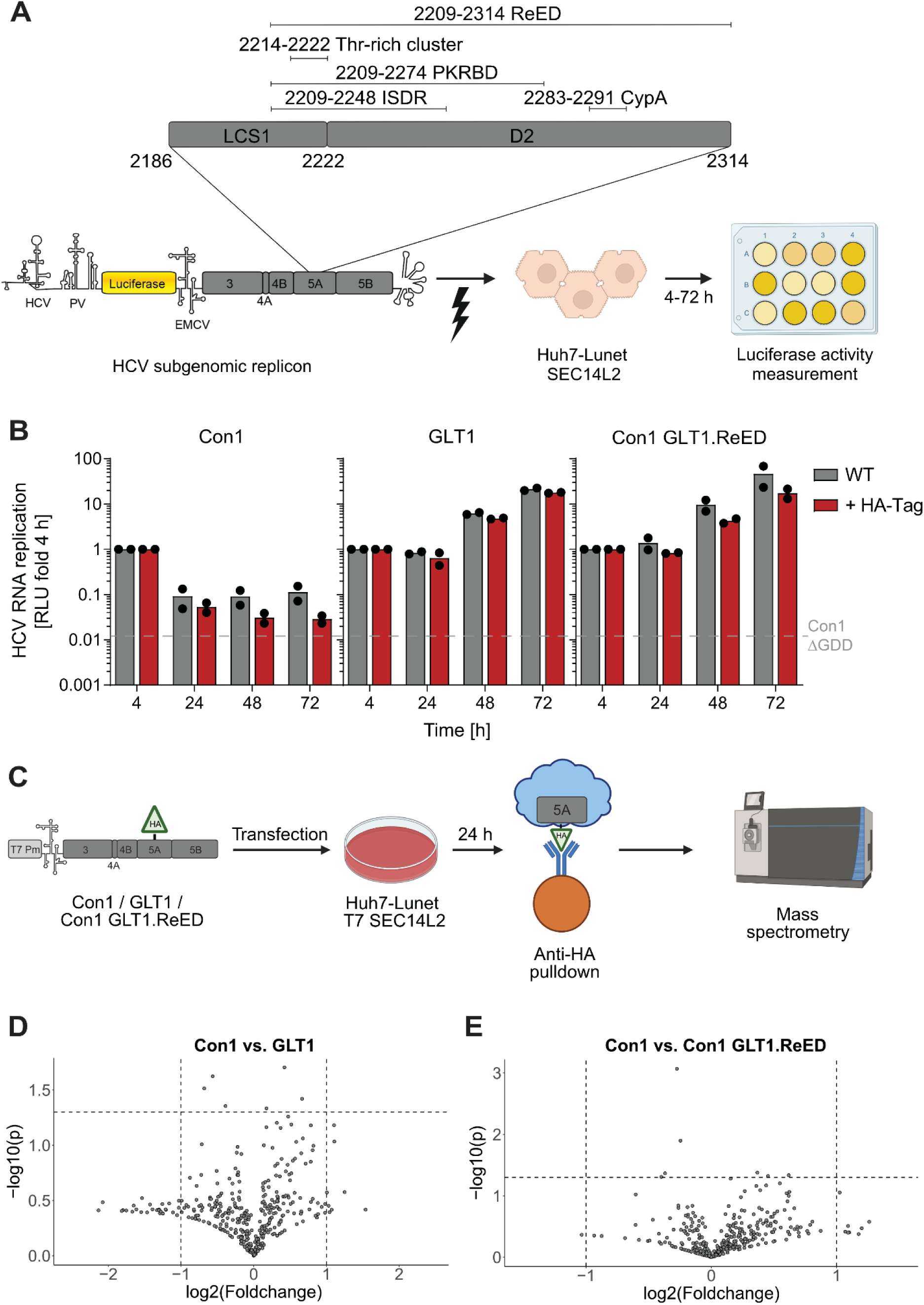
NS5A interactome in the context of different ReEDs. A) Schematic illustrating previously characterised regions in the ReED (21) including a threonine rich cluster crucial for phosphorylation (91), the PKR binding domain (PKRBD) (23), the interferon sensitivity determining region (ISDR) (22) and an interaction site with CypA (24) (upper panel). Schematic illustrating subgenomic replicons (SGRs) harbouring the HCV replicase plus a luciferase reporter and their electroporation into Huh7-Lunet SEC14L2 cells for assessment of HCV replication fitness. B) SGRs of the indicated constructs were electroporated into Huh7-Lunet SEC14L2 cells, luciferase activity in cell lysates (RLU) was quantified as a correlate of RNA replication efficiency at the given time points and normalised to 4 h to account for differences in transfection efficiency. Con1 ΔGDD served as a replication deficient negative control. Data are from two independent biological replicates measured in technical duplicates. Each dot depicts the result of one replicate and the bar indicates the mean of all replicates. C) Schematic illustrating the workflow of T7 polymerase driven protein expression, HA-targeted pulldown and subsequent mass spectrometry. D&E) Volcano plots depicting the foldchange and statistical significance of protein detection between the indicated conditions. Dotted lines represent cutoff values for biological relevance (mean foldchange > 2 or < −2) or statistical significance (p < 0.05). Statistical significance was evaluated by a two-sample t-test corrected for multiple testing through the method of Benjamini-Hochberg. A&C) Schematics were Created in BioRender. Lohmann, V. (2026) https://BioRender.com/zd9aw1i

Mass spectrometry revealed a plethora of NS5A interactors but none of them showed a statistically significant ReED dependent change in interaction (Fig. 1D&E). The correct identification of known NS5A interactors like PI4KA (28), VAPA (29) or NAP1L1 (30), and overall low background binding in samples lacking an HA-tagged construct as compared to the HA-tagged samples (Fig. S1A) confirmed the technical soundness of our analysis. While not statistically significant, a trend towards a lower NS5A-PI4KA interaction was observed for constructs with the GLT1 ReED in line with previous observations of lower PI4P induction by GLT1 (26). Surprisingly, the reported ReED interactor CypA (24) showed similar enrichment in NS5A constructs regardless of the nature of the ReED and the presence of an HA-tag (Fig. S1B). Consequently, we saw a lack of ReED dependent effects on replication inhibition by the CypA inhibitor Alisporivir (Fig. S1C) (31). Overall, these results argued against altered NS5A-host factor interactions as the key mechanism of ReED mediated replication enhancement.

### The ReED mainly has a cis-acting function

Since the interactome analysis did not support an important role of the ReED in mediating interactions with host factors, we focussed on the HCV replicase. It was previously shown that different NS5A functions could be assessed through trans-complementation (32–34). Thus, we co-electroporated a low replicator (Con1) and a high replicator (GLT1) SGR with only the Con1 construct harbouring a luciferase reporter, allowing us to study a potential trans-complementation of the high replicator phenotype by GLT1 (Fig. 2A). However, no trans-complementation was observed since co-electroporation of GLT1 actually reduced Con1 replication (Fig. 2B). To exclude an excessive consumption of cellular resources by GLT1, the experiment was repeated with a replication deficient GLT1 variant only allowing for initial translation but also in this setting co-electroporation did not enhance Con1 replication (Fig. 2C).

**Fig. 2:**
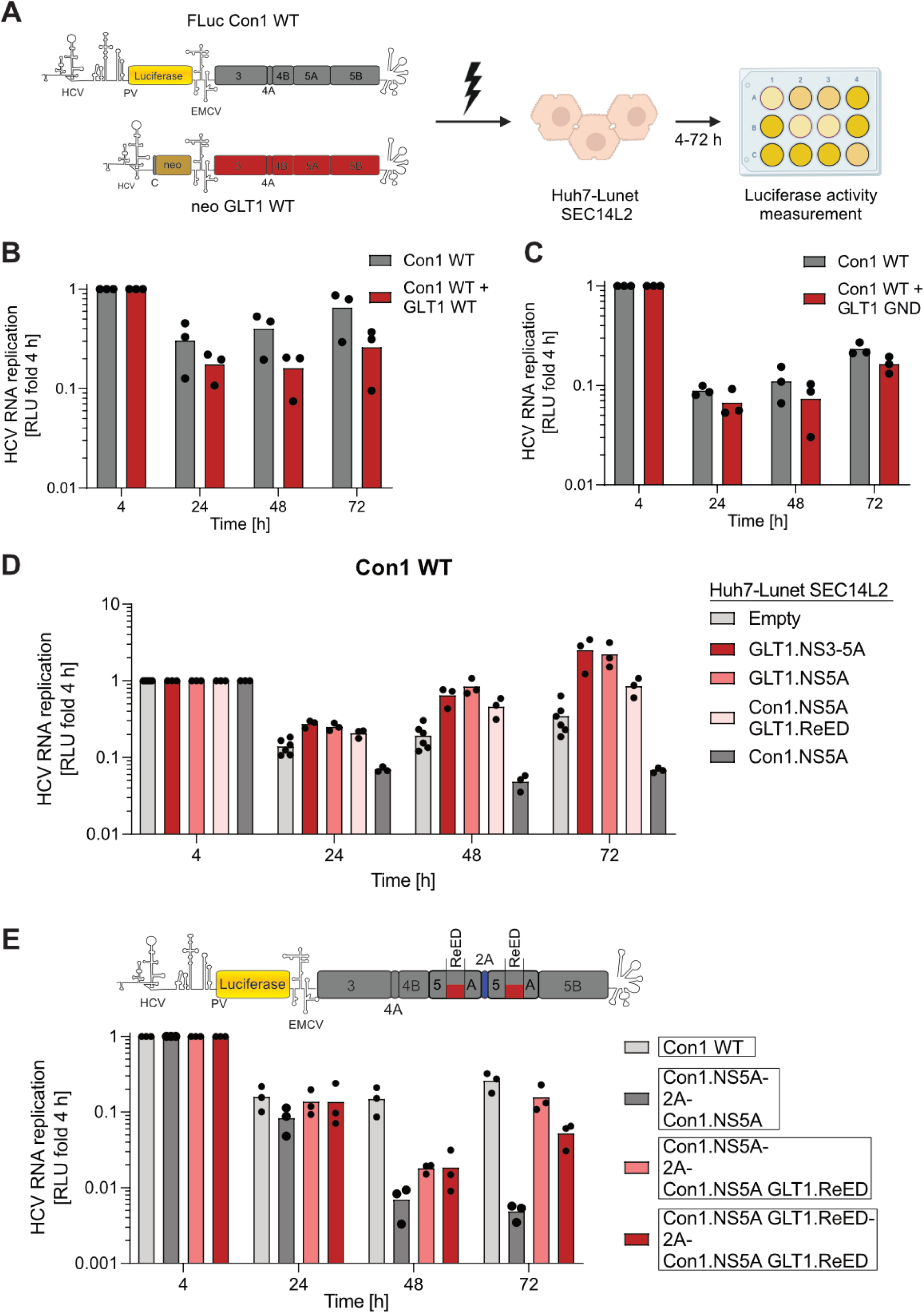
Trans-complementation and ReED dominance. A) Schematic illustrating the co-electroporation of two SGRs with only Con1 containing the luciferase reporter needed for monitoring of replication. Created in BioRender. Lohmann, V. (2026) https://BioRender.com/t48e218 B&C) HCV replication assay was performed as shown in A) with the indicated constructs. Luciferase activity in cell lysates (RLU) was quantified as a correlate of RNA replication efficiency at the given time points and normalised to 4 h to account for differences in transfection efficiency. D) HCV replication assay was performed as shown in Fig. 1A with a Con1 WT SGR. Huh7-Lunet SEC14L2 cells were overexpressing the indicated HCV proteins. Empty refers to a puromycin resistance cassette without additional HCV proteins. E) SGRs with a duplicated NS5A separated by a 2A site allowing for separation of the proteins were used. HCV replication assay was performed as shown in Fig. 1A with the indicated constructs. B-E) Each dot depicts the result of one replicate and the bar indicates the mean of all replicates.

After co-electroporation, production of low and high replicator NS5A variants occurred at the same time but from independent RNAs, potentially leading to compartmentalisation of low and high replicator variants. To avoid such spatial separation, we generated hepatoma cells overexpressing either NS3-5A or NS5A of GLT1 (Fig. S2A). Thereby, the high replicator proteins were already present in the cell upon SGR transfection. NS3-5A was used since previous studies highlighted that trans-complementation of some NS5A functions required the presence of other NS proteins (35). Indeed, overexpression of either GLT1 NS3-5A or GLT1 NS5A increased Con1 replication by roughly 6-fold compared to a control cell line (Fig. 2D). A milder replication enhancing effect on Con1 was observed when overexpressing Con1 NS5A with the GLT1 ReED while Con1 NS5A overexpression decreased replication. When transfecting these cell lines with a high replicator SGR, replication was decreased over 30-fold compared to a control cell line (Fig. S2B).

To further clarify the interplay of low and high replicator ReEDs, we assessed whether a high replicator had a dominant positive effect. To this end, we duplicated NS5A separated by foot-and-mouth disease virus 2A (36) leading to separation of the proteins in the translated SGR (Fig. 2E, upper panel). The constructs with duplicated Con1 NS5A replicated lower than a Con1 WT control (Fig. 2E, lower panel). Nevertheless, the presence of one GLT1 ReED strongly increased replication with no additive effect if both variants harboured the GLT1 ReED. This highlighted that a high replicator ReED indeed has a dominant positive effect on low replicator variants.

Altogether, studies on trans-complementation were inconclusive with co-electroporation arguing for a cis-acting function while protein overexpression prior to electroporation allowed for some trans-complementation moderately stimulating replication of low replicator variants. In a cis-setting with duplicated NS5A, one copy of a high replicator ReED was sufficient to promote replication, arguing for a dominant positive effect of high replicator ReEDs.

### The ReED affects NS5A dimerization

The difficulty of achieving trans-complementation combined with the clear benefit of at least one high replicator ReED copy in the constructs with a duplicated NS5A was in line with the hypothesis that the ReED acted within the HCV replicase. Along these lines, we previously observed that low dose treatment (1-10 pM) with clinically approved NS5A inhibitors phenocopied the replication enhancing effect of a high replicator ReED (21). These inhibitors were proposed to fix NS5A in one dimeric conformation thereby preventing conformational flexibility required for its function (37). The NS5A inhibitor mediated replication enhancement was so far only studied in the context of the Con1 and GLT1 ReED (21). To check whether this phenotype applied to all gt1b ReEDs we tested the effect of 1 pM Pibrentasvir addition shortly after transfection (Fig. S3) on a larger set of previously characterized low and high replicator gt1b ReEDs (21) (Fig. 3A). Indeed, replication was increased up to 50-fold for all tested ReEDs besides the very high replicator FCH3 indicating that low dose NS5A inhibitor mediated replication enhancement can be applied to gt1b in general. The resistance associated mutation Y93H in domain 1 of NS5A was considered to reduce binding of NS5A inhibitors (36, 38) subsequently affecting drug sensitivity for Daclatasvir but not Pibrentasvir treatment in gt1b SGRs (39, 40). In line with this, we could show that the presence of Y93H abrogated the replication enhancing effect of low dose Daclatasvir but not Pibrentasvir treatment (Fig. 3B). This further underlined that the replication enhancing effect of low dose NS5A inhibitor treatment stems from the drug binding to the same site in NS5A as under high antiviral concentrations and not from an off-target effect.

**Fig. 3:**
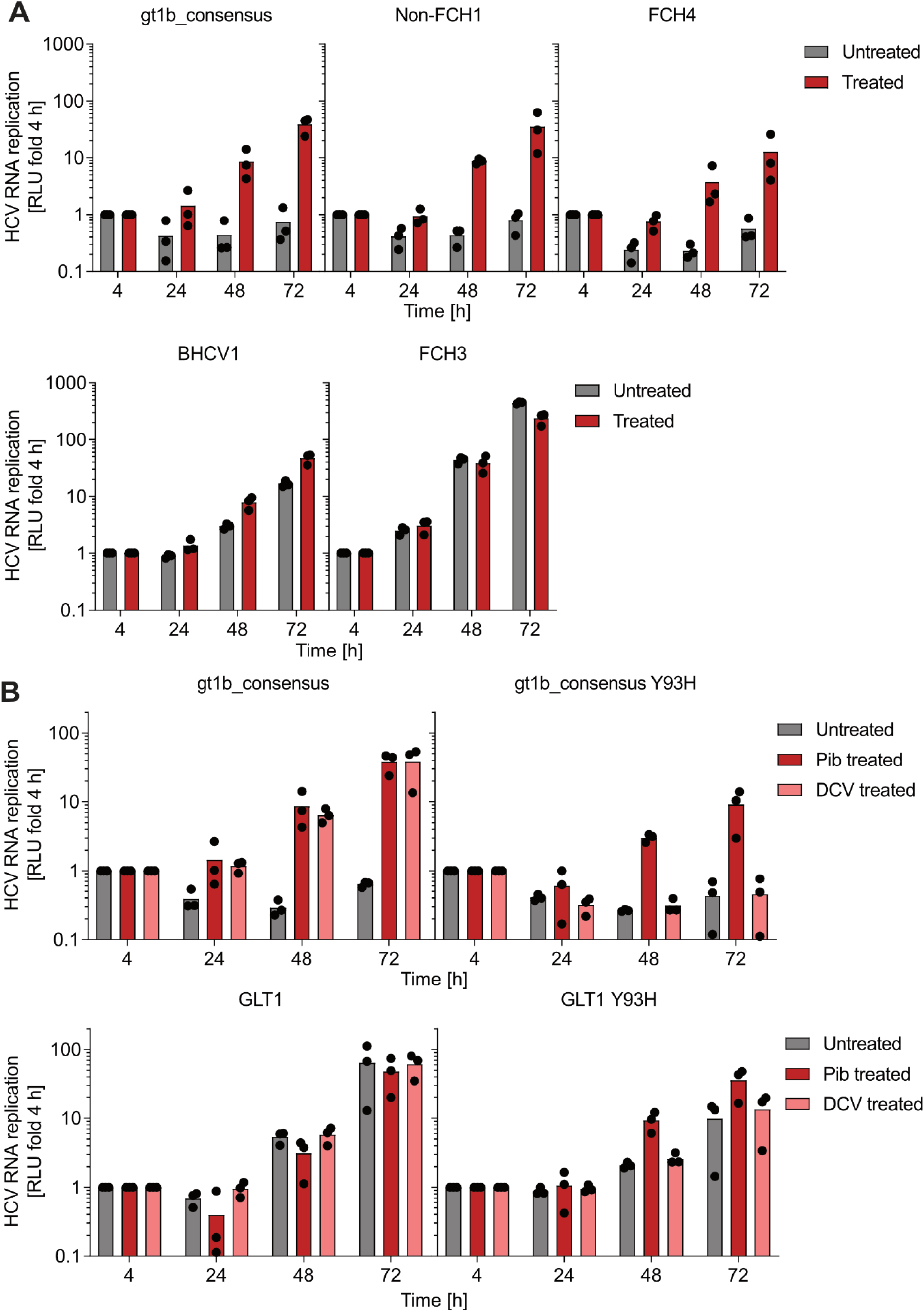
NS5A inhibitor mediated replication enhancement. HCV replication assay was performed with Con1 based SGRs harbouring the indicated ReEDs and NS5A mutations as shown in Fig. 1A. Cells were treated with 1 pM Pibrentasvir (Pib) (A&B) or 10 pM Daclatasvir (DCV) (B) 4 h after transfection. Data are from three independent biological replicates measured in technical duplicates. Each dot depicts the result of one replicate and the line indicates the mean of all replicates.

In order to probe how the ReED might affect the viral replicase, we next performed structural modelling via AlphaFold. AlphaFold2 (41) consistently modelled the ReED with a ∼33-residue segment around positions 2221-2253, i.e. the very N-terminus of D2, as structured into an alpha-helix followed by a two-strand beta sheet, with good confidence statistics (Fig. 4A). In our hands, such isolated structural segments in AlphaFold models generally signal an intrinsically disordered or incomplete segment that folds only upon binding to its partner(s). Indeed, molecular dynamics simulations starting from the folded ReED segment led to a considerable loss of structure, with only part of the alpha helix retained (Fig. 4B). We tried modelling the ReED with NS5B as the most likely putative partner in the replicase (42, 43), but the resulting interaction predictions were unconvincing and poorly scored. As NS5A was shown to form functional homodimers through its structured domain 1 (13, 15), we also tried modelling an interaction of NS5B with two ReED molecules. This also failed but, surprisingly, the same 33-residue segment in the ReED now self-associated into dimers. Indeed, modelling a dimer of this segment on its own yielded a very well-scored prediction (Fig. 4C), that was furthermore stable in molecular dynamics simulations (Fig. 4D). Interestingly, mutations or insertions were often found in or close to this segment in highly replicating strains (Fig. S4A). Similar dimer predictions were obtained for gt1b ReEDs from a variety of gt1b strains (Fig. S4B). We are thus led to an overall structural model of NS5A in which dimers can form through domain 1 as previously reported, but also through the ReED (Fig. 4E).

**Fig. 4:**
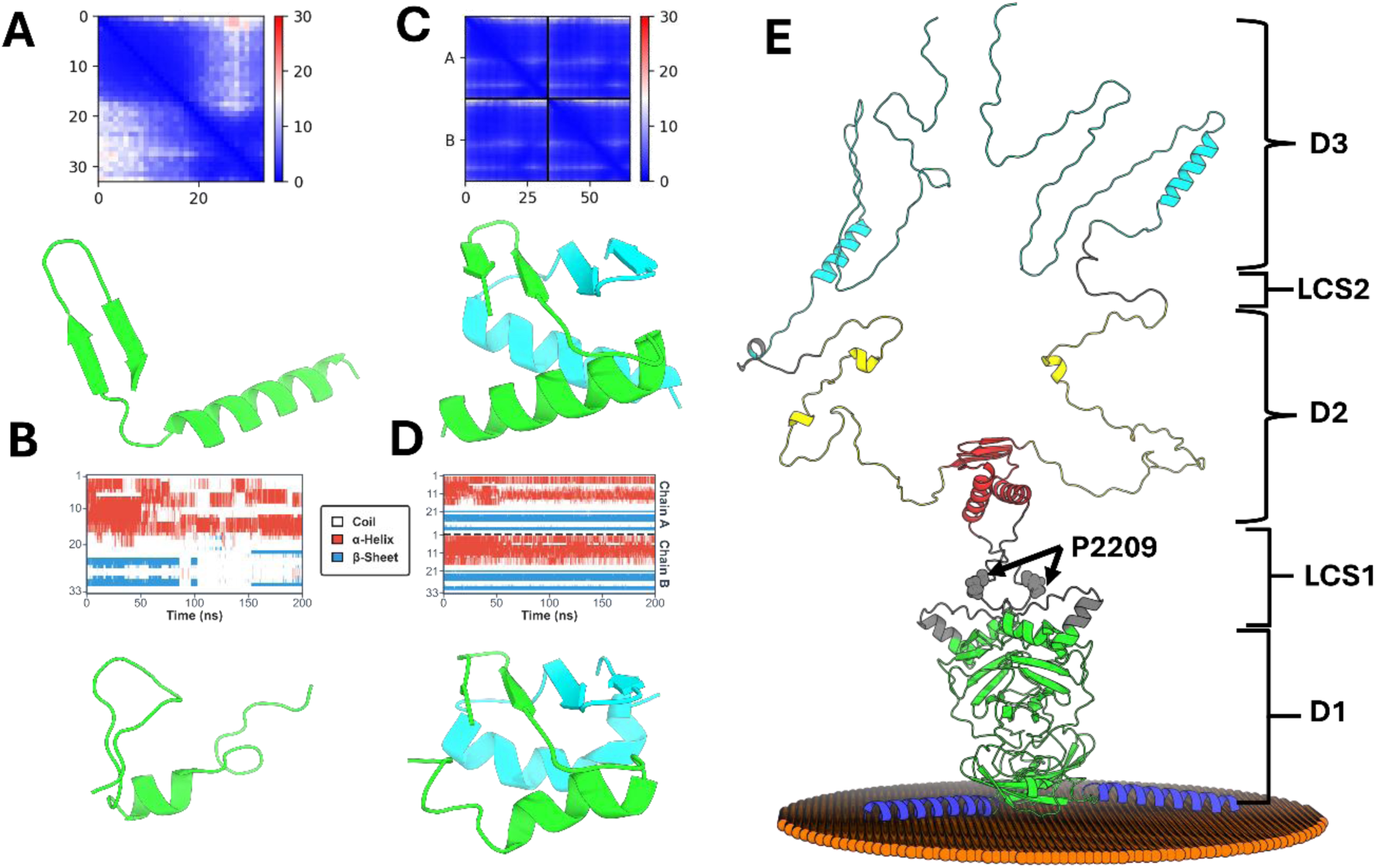
Modelling of a Con1 ReED dimer. A) AlphaFold modelling of an NS5A peptide within the ReED (residues 2221-2253 of the Con1 polyprotein) and B) its molecular dynamics simulation. C) Prediction of a dimer of the same peptide and D) its molecular dynamics simulation. A&C) Top, predicted aligned error (PAE) plots between pairs of residues. PAE is defined as the expected positional error in Ångströms (Å) at one residue if the predicted and actual structures were aligned on the other residue. The scale is from dark blue (0 Å expected error) to dark red (30 Å expected error). Low PAE values off-diagonal indicate high confidence in the 3D structures. Bottom, ribbon representation of the predicted models with the first peptide (chain A) in green and the second peptide (in the dimer) in cyan (chain B). B&D) Top, evolution of the secondary structure contents along the 200-ns simulations (red: residues in helical conformation; Blue: residues in beta-sheet conformation). Bottom, snapshots at the end of the simulations. E) Overall AlphaFold modelling of a Con1 NS5A dimer at a membrane surface (localized by orange spheres). Domain 1 in green with the N-terminal amphipathic helix in blue; LCS1 in grey with the N-terminal residue of the ReED P2209 highlighted as spheres and labelled; Domain 2 in yellow with the putative new dimerization site at its N-terminus in red; LCS2 in grey; Domain 3 in cyan. Boundaries according to (92).

To phenotypically characterise this putative dimerization site and its relevance for replication fitness, we established an NS5A dimerization assay based on split nano luciferase (NanoLuc). Here, constructs allowing for expression of Con1 NS5A harbouring different ReEDs either C-terminally tagged with the small or the large bit were co-transfected into Huh7-Lunet cells. Upon NS5A dimerization, the two bits form a functioning NanoLuc allowing for a quantification of dimerization (Fig. 5A). Employing this method, we found that NS5A with a high replicator ReED showed an on average three times higher luciferase activity than variants with a low replicator ReED (Fig. 5B), with an overall highly significant difference among the groups, suggesting a crucial role of ReED mediated NS5A dimerization in the regulation of HCV replication fitness. An NS5A variant with ReED deletion indeed showed strongly decreased dimerization signals, only sixfold higher than the eGFP negative control (Fig. 5B), which were likely mediated by D1 dimerization. Therefore, the majority of the signal detected in this assay was based on ReED mediated dimerization, potentially due to placement of the NanoLuc fragments on the C-terminus of NS5A which is spatially closer to the ReED than to D1. ReED-mediated dimerization enhancement was confirmed by comparing NS5A with the gt1b consensus ReED with a mutant harbouring only the P2209L mutation, which was previously shown to enhance replication fitness (21) and indeed stimulated dimerization (Fig. 5C). In contrast, when applying 1 pM Pibrentasvir to the NS5A dimerization assay, which phenocopied the ReED mediated replication enhancement, no effect was observed even at high drug doses (Fig. 5D). This might be due to the drug binding to D1 and thus likely affecting D1 mediated dimerization, which was only weakly detected with this assay.

**Fig. 5:**
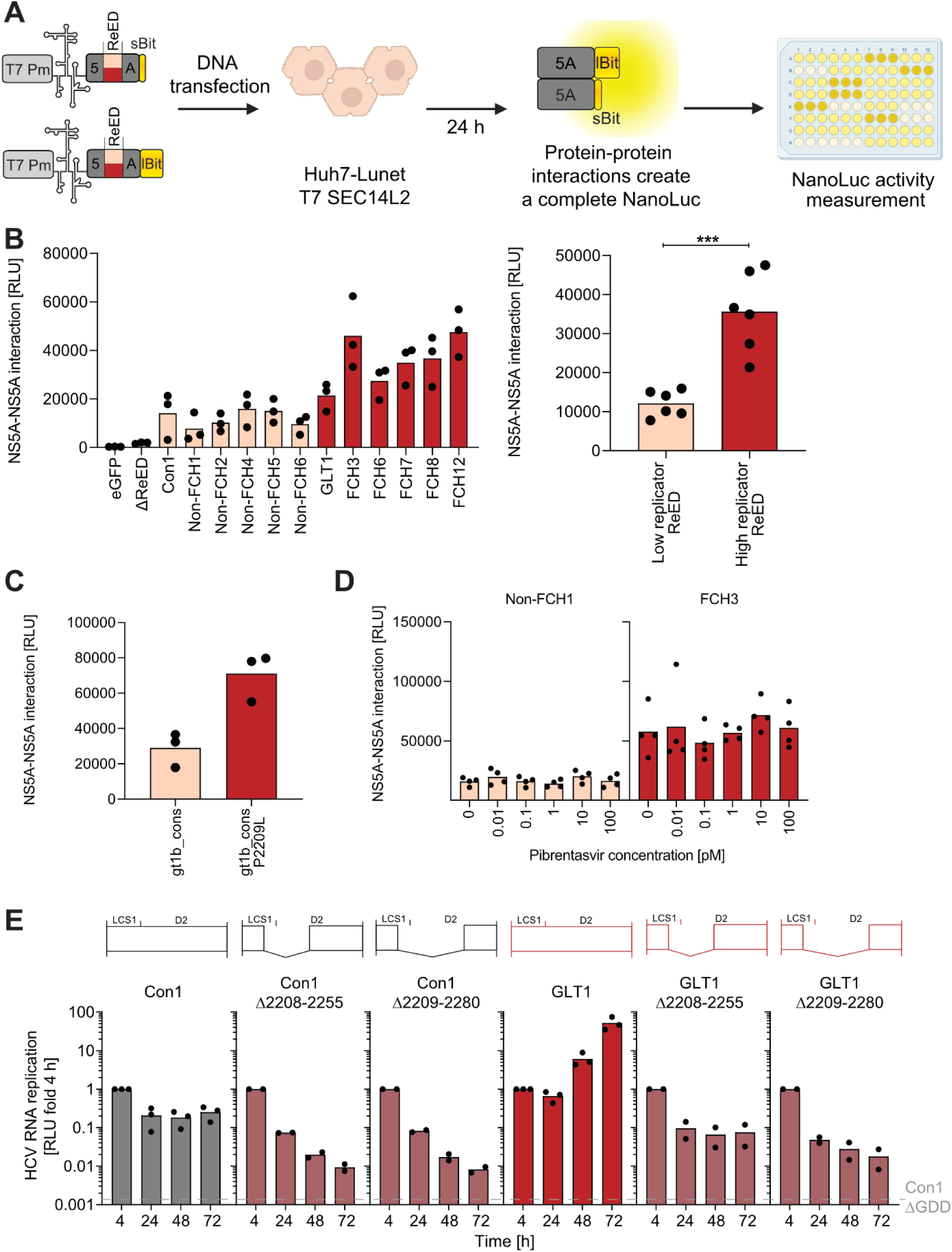
ReED dependent effects on NS5A dimerization. A) Schematic of the NS5A dimerization assay, co-transfecting two constructs harbouring Con1 NS5A based ReED chimeras tagged with either the sBit or the lBit into Huh7-Lunet T7 SEC14L2 cells. Protein expression was mediated by T7 polymerase. Cells were lysed after 24 h and the NanoLuc signal created through NS5A dimerization was measured. Created in BioRender. Lohmann, V. (2026) https://BioRender.com/qofj4u0 B-D) NanoLuc activity in cell lysates (RLU) was quantified as a correlate of NS5A homodimerization using Con1 NS5A constructs harbouring the indicated ReEDs. Constructs shown with an apricot-coloured bar contained ReEDs previously shown to induce low replication levels while constructs shown with a red bar contained ReEDs previously shown to induce high replication levels (21). Interaction of Con1 NS5A lacking the ReED (ΔReED) or lBit tagged eGFP with sBit tagged Con1 NS5A served as negative controls. B) The right panel is an aggregation of the mean values per construct form the left panel based on the presence of a low or high replicator ReED. Statistical significance was determined with a two-sided Student’s t-test. *** = p < 0.001. E) HCV replication assay was performed as shown in Fig. 1A with Con1 or GLT1 based constructs harbouring the indicated deletions in NS5A. Data are from two or three independent biological replicates measured in technical duplicates. Each dot depicts the result of one replicate and the line indicates the mean of all replicates.

Lastly, we wanted to assess whether the ReED was essential for genome replication. Therefore, we created SGRs harbouring a deletion from the start of the ReED until the CypA binding site or a deletion of the ReED N-terminus which was previously identified to confer cell culture adaptation in a selectable replicon (8). Nevertheless, both deletions led to an at least 20-fold reduction of replication in both Con1 and GLT1 (Fig. 5E), highlighting that the ReED is needed for efficient HCV replication. All in all, the ReED appears to contain a so far overlooked, essential NS5A dimerization site with increased NS5A dimerization correlating with high replication fitness. Treatment with NS5A inhibitors potentially mediating NS5A dimerization via D1 phenocopied the effect of high replicator ReEDs.

### The ReED depends on HCV helicase and polymerase

While the correlation of NS5A dimerization strength and replication fitness was quite striking, it remained unclear how the stronger dimerization alters NS5A function. A previous publication highlighted an interaction between NS5A D2 which is part of the ReED and the RNA dependent RNA polymerase NS5B which downregulated NS5B activity (43). To analyse whether the ReED affects NS5B activity, we established an RdRp activity assay (44) based on purified recombinant Con1 NS5B (45). Additionally, we purified recombinant Con1 LCS1D2 protein fragments containing the ReED of isolates Con1, GLT1 and FCH3 (Fig. 6A). The latter was the ReED leading to highest replication levels in our previous study (21). When adding low amounts of ReED not exceeding the molarity of NS5B in the RdRp activity assay, a ∼40% enhancement of RdRp activity was observed for all ReEDs (Fig. 6B). When further increasing the ReED concentration exceeding NS5B’s by up to 40-fold, a decrease in RdRp activity of 50% was observed for Con1 while for the high replicator ReEDs GLT1 and FCH3 a 90% decrease was observed compared to an untreated control. These results were counterintuitive since one would expect a higher RdRp activity to be associated with higher replication fitness; however, size exclusion chromatography argued for the majority of purified LCS1D2 protein fragments not dimerizing (Fig. 6C). A small second peak in the variants containing the GLT1 or FCH3 ReED could indicate that a fraction of molecules dimerized with stronger signal for the high replicator ReEDs being in line with our findings so far. Thus, working with purified protein fragments might be a too reductionistic system eliminating key components like membranes, other NS5A domains, other replicase proteins or post translational modifications to just name a few factors precluding dimerization and thus functionality of the ReED.

**Fig. 6:**
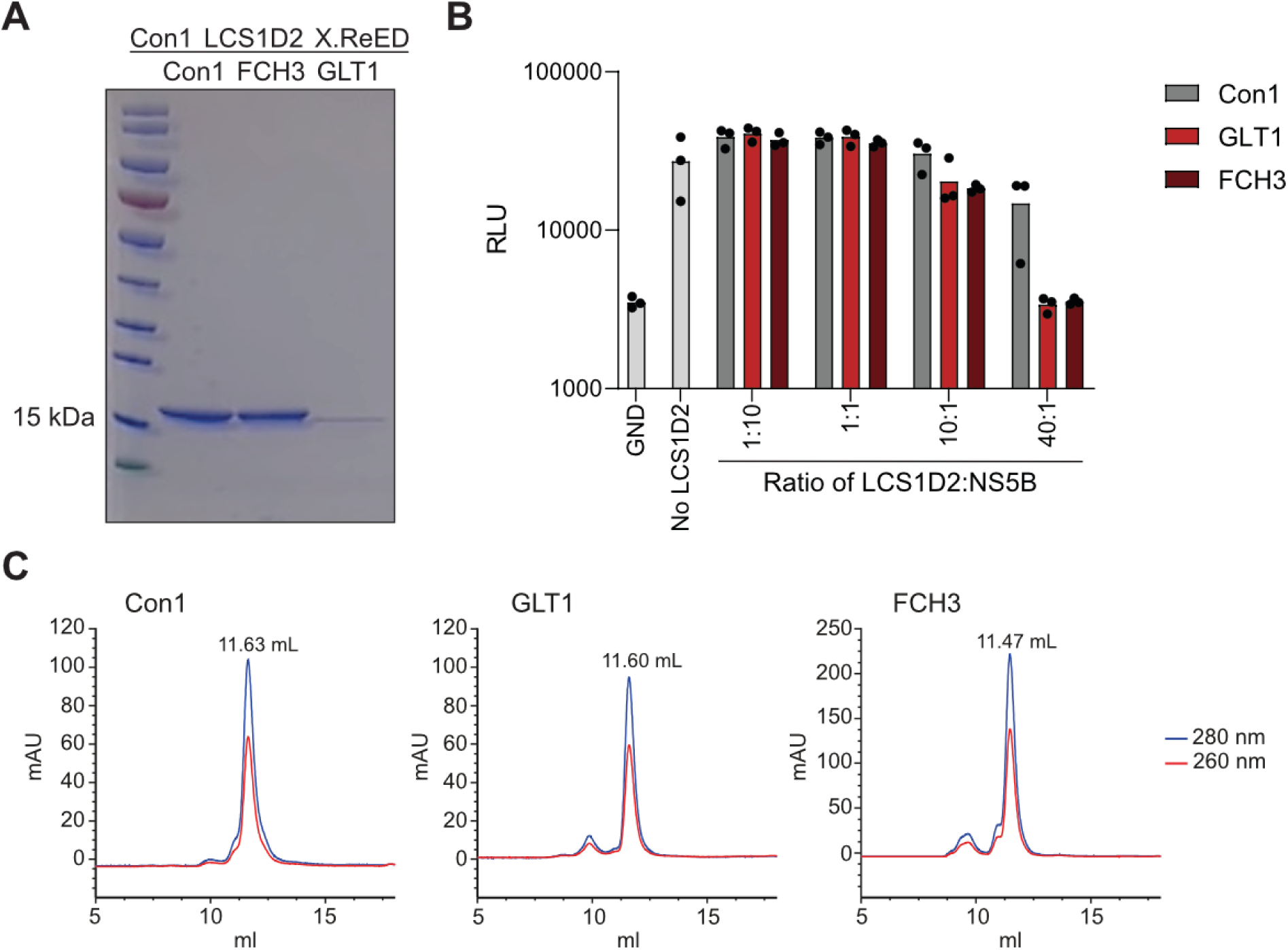
The effect of purified LCS1D2s on RdRp activity. A) Con1 LCS1D2 harboring the indicated ReED and a C-terminal 6x-His tag was expressed in *E. coli* BL21 rosetta. Coomassie stained SDS-PAGE of the purified product. B) PicoGreen fluorescence intensity was measured as a correlate of RNA polymerization by Con1 NS5B. Purified LCS1D2s with the indicated ReED was added in the indicated ratios. JFH1 NS5B GND served as an RNA polymerization deficient negative control. C) The recombinant LCS1D2 proteins with a Con1, GLT1 or FCH3 ReED were analyzed using size-exclusion chromatography (Superdex 75 Increase 10/300 GL). Protein elution was monitored by measuring the absorption at 280 nm and 260 nm (blue and red curves, respectively). The elution volumes of the LCS1D2 proteins are indicated above the chromatograms (mAU: milli-absorbance units).

Since working with purified proteins seemed suboptimal, we focussed on the gt2a isolate JFH1, which is the only non-gt1b HCV WT isolate with very high replication fitness in cell culture, stemming from a fulminant hepatitis patient (7, 45, 46). To study the ReED in gt2a, we further focussed on JFH2, another fulminant hepatitis patient derived isolate (47) and on J6, isolated from an acute phase chimpanzee infected with patient serum from an asymptomatic carrier (48, 49), representing a low replicator variant (46). Interestingly, the ReED of J6 was quite similar to the gt2a consensus while JFH1 and JFH2 had ReEDs differing substantially from the consensus (Fig. 7A), a pattern that was associated with elevated replication fitness in other gts (21). Indeed, J6 with its few ReED mutations showed a low replicator phenotype similar to the gt2a consensus ReED, as expected (Fig. 7B, S5A). Introducing the highly altered JFH2 ReED into J6 induced an over 100-fold increase in genome replication fitness (Fig. 7B). However, the JFH1 ReED actually reduced replication to the level of a replication deficient negative control. This was surprising considering the reported high replicator phenotype of JFH1 (7). Thus, we introduced the J6 and JFH2 ReED into JFH1. Here, replication reached very high levels, 10,000-fold higher than J6, but none of the ReEDs showed any meaningful effect on replication fitness (Fig. 7B), suggesting that JFH1 achieved high replication fitness independent of the ReED.

**Fig. 7:**
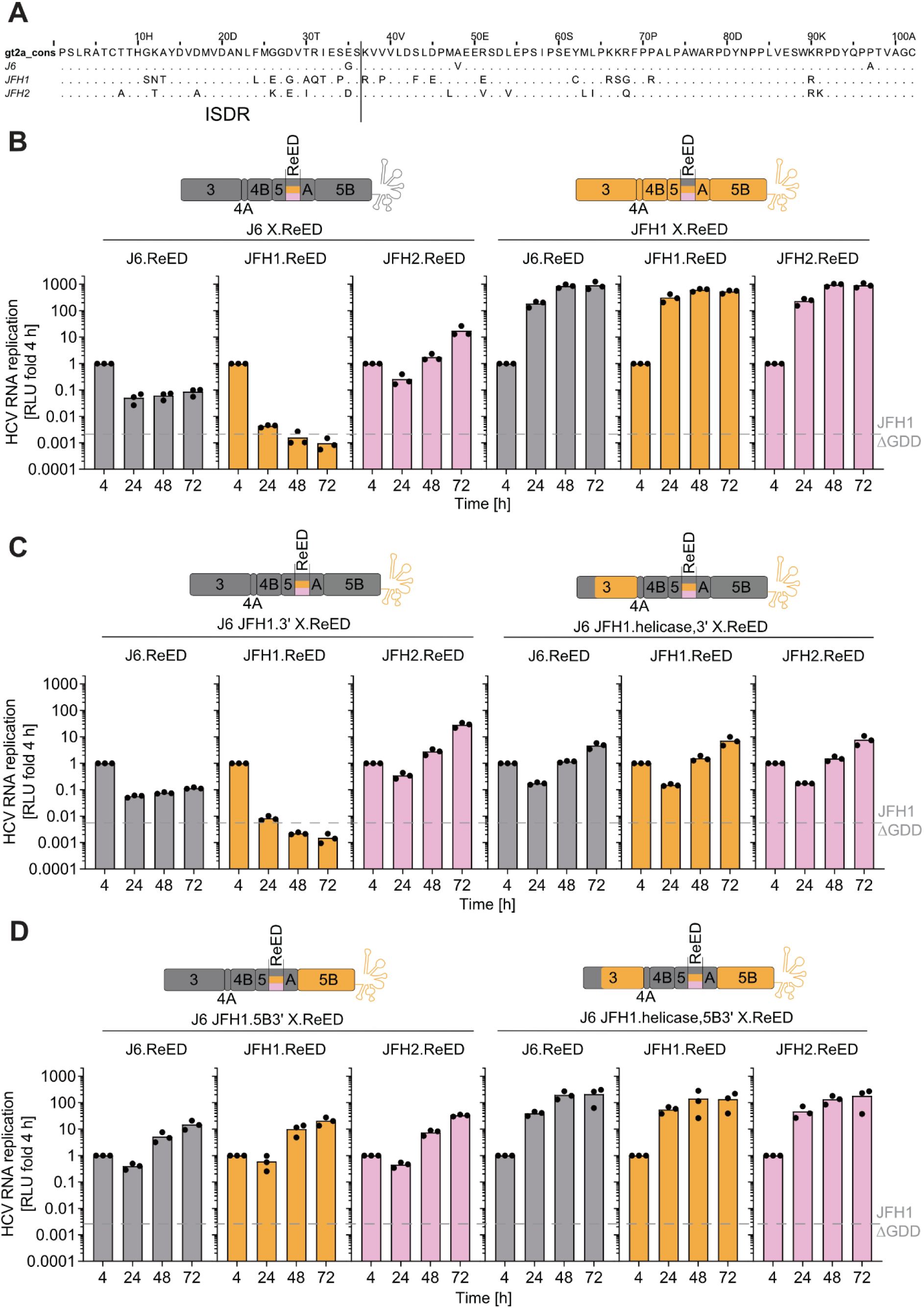
Determinants of ReED resistance in the gt2a high replicator JFH1. A) ReED amino acid alignment between J6, JFH1, JFH2 and the gt2a consensus, dots indicate an amino acid being identical to the gt2a consensus. B-D) HCV replication assay was performed as shown in Fig. 1A. Schematics of the used constructs are shown in the upper panels: J6 or JFH1 with the indicated ReEDs (B), J6 with the JFH1 3’UTR, the indicated ReED with and without the JFH1 NS3 helicase (C) or J6 with the JFH1 3’UTR, the indicated ReED and JFH1 NS5B polymerase with or without JFH1 NS3 helicase (D). JFH1 ΔGDD served as a replication deficient negative control. Data are from three independent biological replicates measured in technical duplicates. Each dot depicts the result of one replicate and the bar indicates the mean of all replicates.

Previous studies revealed that the enzymatic activities of NS3 helicase and the NS5B polymerase in combination with the 3’UTR were central determinants of the outstanding replication efficiency of JFH1 (46). Thus, we aimed at analyzing whether these factors could transfer the ReED resistance of JFH1 to J6 based SGRs. When introducing only the 3’UTR of JFH1 into J6, the resulting construct remained ReED sensitive (Fig. 7C). Indeed, when introducing either JFH1 helicase and 3’UTR or JFH1 polymerase and 3’UTR into the ReED sensitive J6 isolate, the resulting chimera became ReED resistant (Fig. 7C&D). The same effect with an overall higher replication fitness was achieved by a combined transfer of JFH1 helicase, polymerase and 3’UTR into J6 (Fig. 7D). Similarly, transferring just the JFH1 helicase but not the JFH1 3’UTR into J6 resulted in a ReED resistant construct with low replication fitness (Fig. S5B). This showed that the ReED’s replication enhancing capabilities are affected by determinants in helicase and polymerase implying that the ReED is acting directly on the HCV replicase.

To further substantiate the role of helicase and polymerase in ReED resistance, we aimed to render JFH1 based SGRs ReED sensitive through introduction of homologous J6 regions. Transfer of J6 helicase or polymerase individually did not alter ReED sensitivity (Fig. 8A). Only when transferring both components from J6 into a JFH1 SGR, ReED sensitivity was also transferred independent of the 3’UTR present in the construct (Fig. 8B). This confirmed that ReED sensitivity required the presence of helicase and polymerase of a low replicator isolate. Importantly, the level of replication correlated with NS5A dimerization for J6 based chimeric constructs (Fig. 8C), similar to gt1b. This effect was less pronounced in JFH1 based chimeras highlighting its independence of the ReED regulatory function and indicating that ReED dimerization might depend on additional determinants within NS5A (Fig. 8D). Overall, these experiments confirmed the correlation of stronger NS5A dimerization and replication fitness beyond gt1b and identified determinants in helicase and polymerase regulated by the ReED. Based on these data, we propose the following model: NS5A usually has an inhibitory function on helicase and polymerase, stalling the replicase likely prior to initiation of RNA synthesis. Through ReED mediated dimerization this inhibitory function is released, and genome replication is initiated. In high replicators, the ReED mediated dimerization is stronger, thus the inhibitory function is released more frequently or earlier, and RNA replication occurs at a higher rate (Fig. 9).

**Fig. 8:**
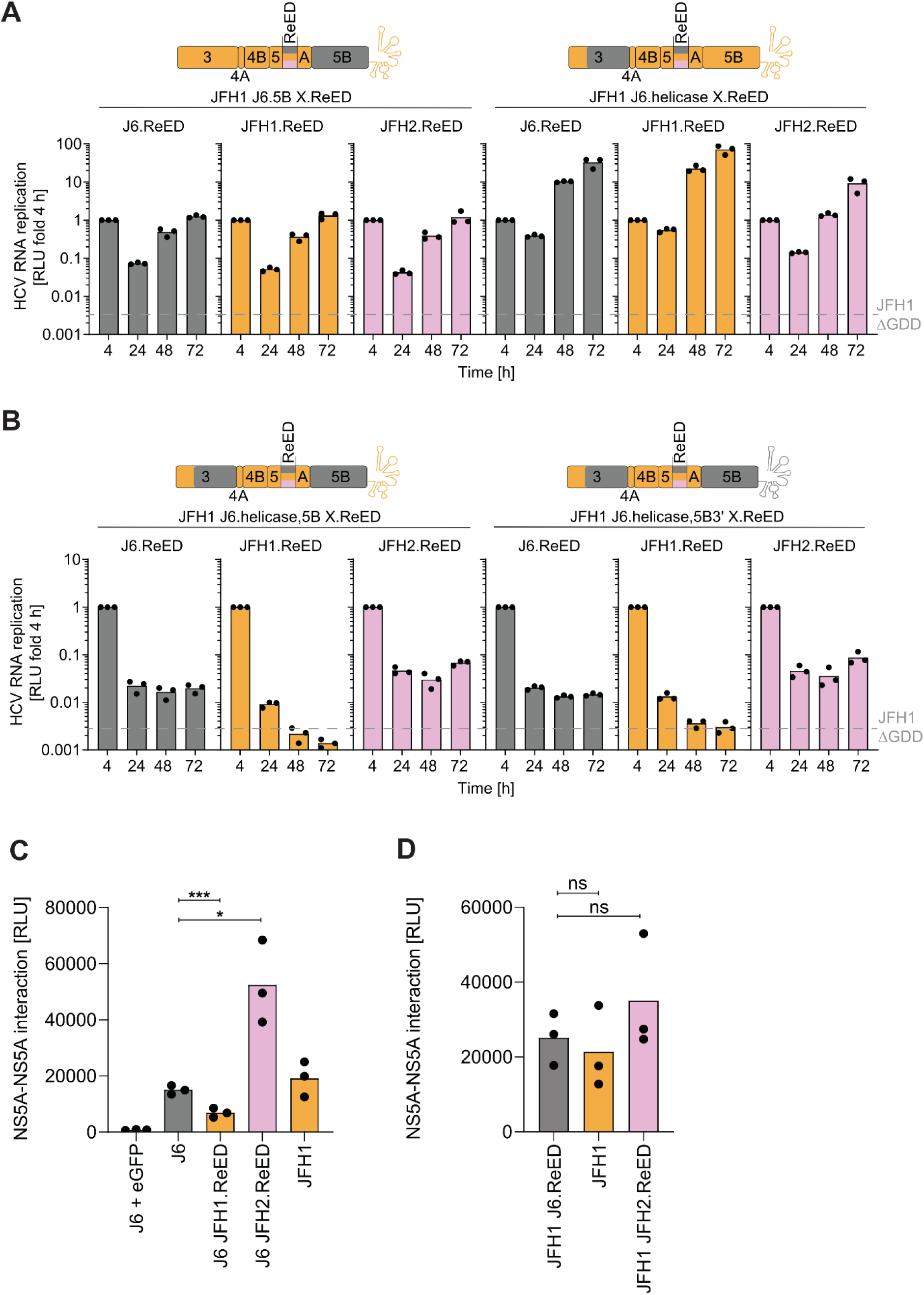
Determinants of ReED sensitivity in the gt2a low replicator J6 and the connection to NS5A dimerization. A&B) HCV replication assay was performed as shown in Fig. 1A. Schematics of the used constructs are shown in the upper panels: JFH1 with the indicated ReED and either the J6 NS3 helicase or NS5B polymerase (A) or JFH1 with J6 NS3 helicase + NS5B polymerase, the indicated ReED with or without J6 3’UTR (B). JFH1 ΔGDD served as a replication deficient negative control. C&D) NS5A dimerization assay was performed as shown in Fig. 5A. NanoLuc activity in cell lysates (RLU) was quantified as a correlate of NS5A dimerization. Interaction of lBit tagged eGFP with sBit tagged J6 NS5A served as a negative control. A-D) Data are from three independent biological replicates measured in technical duplicates (A&B) or triplicates (C&D). Each dot depicts the result of one replicate and the bar indicates the mean of all replicates. Statistical significance was determined with a two-sided Student’s t-test. ns = p > 0.05, * = p < 0.05, *** = p < 0.001.

**Fig. 9:**
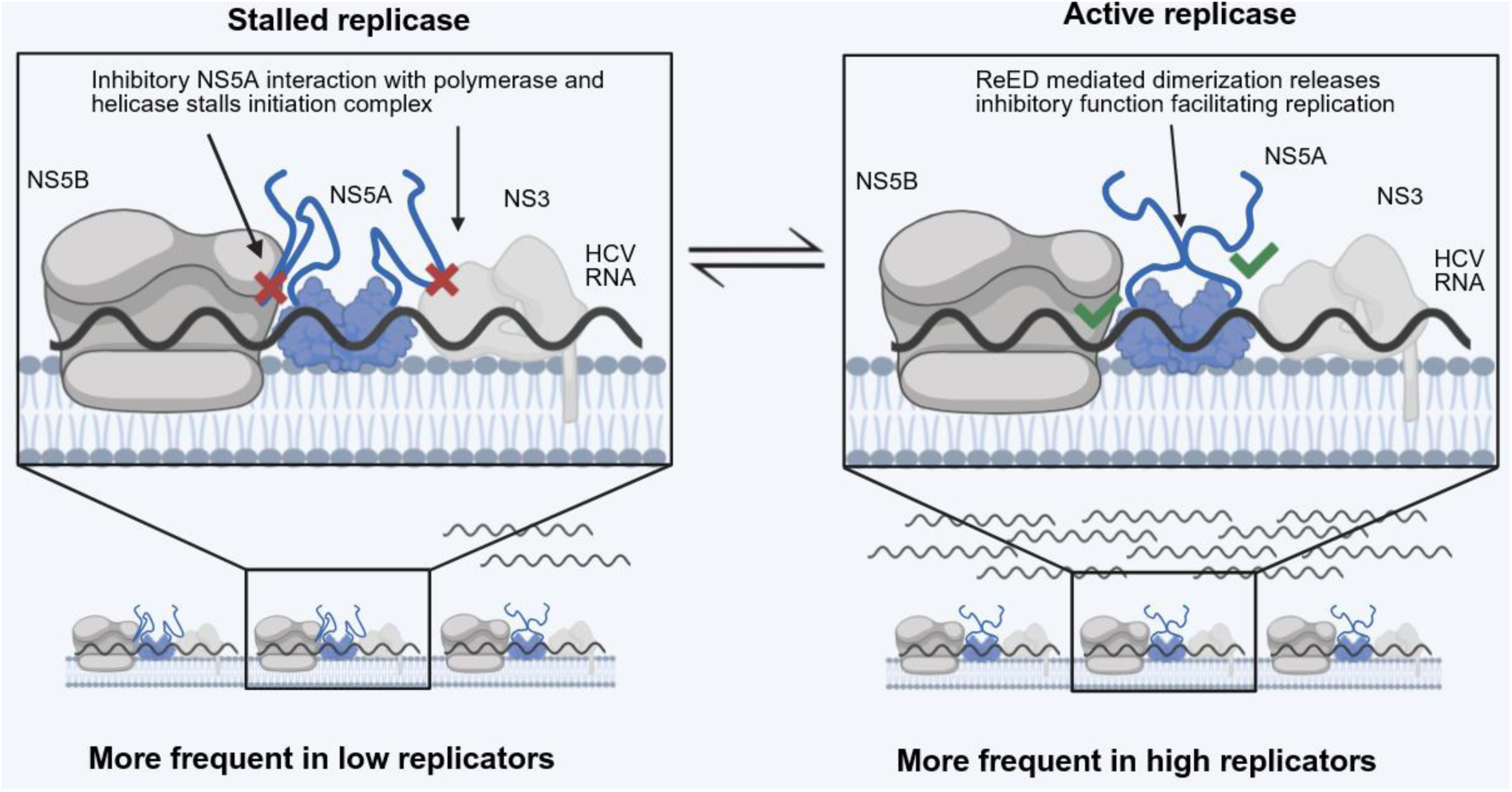
Model of how the ReED regulates HCV genome replication fitness. We assume that NS5A has a negative regulatory function on NS3 helicase and NS5B polymerase. ReED mediated dimerization releases this inhibitory interaction allowing for initiation and subsequent genome replication. Since a high replicator ReED has a higher tendency for dimerization the inhibitory regulation of NS5A is released more often thus initiation and replication occur at a higher rate. Schematic was created in BioRender. Lohmann, V. (2026) https://BioRender.com/2ck2s3u

## Discussion

In the present study, we identified a so far unrecognized NS5A dimerization site within the ReED where mutations appeared to enhance dimerization correlating with elevated replication fitness. The altered NS5A dimerization did not significantly change interactions with host factors but a strong dependence of the replication enhancing effect of the ReED on helicase and polymerase was observed in gt2a. This strongly argues for the ReED acting directly within the HCV replicase.

In general, NS5A dimerization was shown to be necessary for HCV replication (14). Formation of oligomeric NS5A structures involving multiple binding interfaces was also suggested (13–15). Usually, NS5A was considered to dimerize through D1 with a variety of structures proposed (13, 15, 50, 51), accordingly one study found no dimerization if just D2D3 was expressed (18). This is in line with the purified LCS1D2 peptides tested here. Nevertheless, our AlphaFold modelling argued for another, perhaps secondary NS5A dimerization site in the N-terminus of D2 and thus the ReED. Key limitations of AlphaFold include i) a lack of correlation between quality indicators and binding affinity in a complex and ii) models that are insensitive to the effects of point mutations (52). Because of i), a highly scored complex, such as our prediction of a Con1 2221-2253 dimer, is likely accurate but may be of low or high affinity. Our molecular dynamics simulations do show that once formed, this dimer seems to be fairly stable. But it is not feasible to assess how fast it would form, especially in the context of full-length NS5A. While a handful of scattered mutations may be important for dimer formation and/or stability, because of ii) we cannot expect to find obvious differences in dimer structures between predictions of closely related sequences. Indeed, we find a structuration into a dimer around the same helix-strand-strand motif for all sequences of gt1b high and low replicators tested. Still, we may speculate that key replication enhancing mutations in high replicator ReEDs substituting P2209 (21) may lead to different preferred orientations of D2 relative to D1, facilitating encounter of two ReEDs. Similarly, it may be worth noting that in high replicators that have kept the consensus P2209, structural predictions tend to add extra helices to the central motif, especially in strains with clustered substitutions (*e.g.* BHCV1) or with an insertion (*e.g.* FCH5). This may help and/or stabilize folding and dimerization.

In our previous study, we made the surprising observation that low dose NS5A inhibitor treatment enhanced HCV replication (21). Here, we could confirm this effect for a variety of ReEDs of gt1b, where the effect appears to be much stronger than in gt1a or gt3a (53). The replication enhancing effect was abrogated by the presence of the resistance mutation Y93H in a drug specific manner indicating that DAA induced replication enhancement is achieved through the same mechanism that has antiviral effects in larger drug concentrations. The details of the interaction of NS5A inhibitors with their target protein remains a matter of debate with asymmetric and symmetric binding to NS5A dimers proposed (37, 54–56). Especially models of symmetrical binding to the dimer interface fostered speculation that NS5A inhibitors act by fixing NS5A in distinct dimer conformations (37). A low amount of NS5A inhibitor might therefore enforce a subpopulation of NS5A dimers to adopt a conformation that leads to efficient genome replication further arguing for a connection between replication fitness and NS5A dimerization. This might be mediated by a release of (inhibitory) NS5A interaction with RNA, as suggested by a previous study (57). Structural biology studies using cryo-EM or even NMR could clarify the precise structural differences of NS5A dimers with a low or high replicator ReED in presence and absence of low dose NS5A inhibitor treatment. For cells transfected with cell culture adapted Con1, it was shown that around 1 million copies of each non-structural protein were present in a cell after 48 h (58). The replication enhancing effect of Pibrentasvir was achieved at a concentration of 1 pM. Even under the unrealistic assumption of complete uptake just ∼3500 drug molecules per cell would have been available in our experimental conditions, implying a strong excess of NS5A over the drug. On the other hand, compared to the 1 million NS protein copies, only 40 active replication complexes were detected (58). Thus, NS5A inhibitors might specifically bind NS5A dimers involved in active genome replication implying that formation of a distinct NS5A dimer conformation could be a key regulator of genome replication. Overall, this underlines how the replication enhancing effect of low dose NS5A inhibitor treatment is achieved by targeting a minor subpopulation of NS5A molecules. Conversely, the altered dimerization mediated by high replicator ReEDs might just affect the small fraction of NS5A molecules involved in active replication. However, both options are consistent with our model that NS5A dimerization releases an inhibitory function of NS5A on the replicase by different molecular mechanisms.

The changes in NS5A dimerization likely affect the activity of NS3 helicase and NS5B polymerase as revealed by the gt2a chimeras. Both helicase and polymerase were previously identified as the drivers of high replication fitness of JFH1 (46, 59). More detailed studies on recombinant proteins revealed that the JFH1 helicase had a higher RNA binding affinity and accomplished more immediate unwinding (60). In the same vein, NS5B of JFH1 was also shown to have a higher polymerase activity due to more efficient initiation (61). Since JFH1 helicase and polymerase mediated ReED resistance, it appears that the intrinsically elevated enzymatic activity of both proteins might be key in withstanding the regulatory effects of the ReED further arguing for the ReED usually having a negative regulatory function on helicase and polymerase. For all these factors, a complex network of interactions was described in literature, especially the interplay of NS5A and NS5B. Several studies postulated an interaction of the N-terminus of D2 with NS5B for both gt1b and gt2a (42, 43, 62, 63) with one NMR study claiming interaction sites with NS5B all over the ReED (64). Within NS5B, several sites crucial for NS5A interaction were described (43, 65). Mutations that ablate this interaction rendered SGRs replication incompetent highlighting their importance (66). How binding of NS5A to NS5B affects RdRp activity is not fully understood yet. One study suggested that NS5A D2 bound to residues in the RNA exit channel of NS5B and decreased NS5B-RNA binding which could decrease RNA synthesis rates (43). Along these lines, NS5B conformational changes with subsequent reduction in RNA binding were similar upon binding of D2 or the allosteric NS5B inhibitor filibuvir (63). In vitro RdRp activity assays often described low amounts of NS5A as stimulatory for NS5B activity (42, 67) with higher concentrations showing an inhibitory effect (42, 43). Our data for the RdRp assay was partly in line with literature, also observing a slight boost at low ReED concentrations and a drop of NS5B activity with higher ReED abundance. However, for meaningful assessment of differing roles of low vs. high replicator ReEDs crucial factors like membranes, other replicase components or the ability of our LCS1D2 protein fragments to dimerize efficiently were missing. Deeper insights could be gained by purification of authentic replication complexes allowing for structural and biochemical characterisations. All in all, literature supports the concept of the ReED having a negative regulatory function on NS5B.

In addition to the studies on D2-NS5B interaction, in vitro studies also pointed out that helicase and polymerase could modulate each other’s enzymatic activity (68, 69) and that the helicase bound to the UTRs of the HCV genome (70, 71). Since all factors showing an impact on replication fitness discussed here (NS3, NS5A, NS5B, 3’UTR) were shown to affect initiation of replication (72), it is tempting to speculate that the ReED exerts its regulatory function on this specific process. Because NS5A is an RNA binding protein with D1D2 being required for RNA binding (18) future studies should also examine potential ReED dependent differences in NS5A-RNA interaction. Initiation of RNA replication is also a key process during the synthesis of negative strand RNA serving as a template for the synthesis of the plus strand HCV genome. The ReED might also interfere here since one in vitro study on RdRp activity highlighted the stimulatory effect of NS5A especially on negative strand synthesis (67) and the NS3 helicase was recently shown to be crucial for negative strand synthesis (71). In conclusion, all factors shown to affect ReED activity also appear to be involved in the initiation of RNA synthesis which is regulated by a tight interplay of all factors.

We assume that ReED mediated negative regulation of helicase and polymerase is widely independent from the role of NS5A on the formation of the membranous web (73). Here, recruitment and activation of PI4KA has been shown to be critically involved in promoting phosphatidylinositol-4-phosphate (PI4P) production (28, 74), which has an essential concentration dependent function in shaping the size and morphology of DMVs (11), which is furthermore critically affected by NS5A inhibitors (36, 75). Interestingly, a very recent study demonstrated that direct binding of PI4P to NS5A D1-dimers might induce a conformational switch contributing to membrane reorganization and membranous web formation (76). The regulatory sequences in NS5A governing PI4KA activation, involving the PI4KA functional interaction site and a distinct phosphorylation pattern, are located at the C-terminus of D1 and in LCS1 (11, 35), directly preceding the ReED. It is tempting to speculate that the inhibitory function of the ReED might prevent initiation of RNA synthesis until the formation of replication organelles has occurred, which might further provide the trigger for D2 dimerization. ReED mediated NS5A dimerization might therefore be another step in a series of temporal regulatory events governed by distinct dimer conformations of NS5A.

Overall, data from large sequencing studies suggest low replication fitness being optimal for HCV in achieving persistent long-term infection with slowly progressing pathogenesis (21). High replicators were mainly associated with severe courses of disease in liver transplant patients (21) or as shown here through JFH1 and JFH2 with fulminant hepatitis. Thus, the ReED exerting a negative regulatory function on replication fitness appears to be a key ability of HCV for persistence implying high replicator ReEDs being a rare loss of this regulation only benefiting the virus in extreme conditions like immunosuppression. Taken together, we propose that the tight interplay of NS proteins and viral RNA is modulated by the ReED affecting initiation of RNA synthesis. Especially the literature on NS5A-NS5B interaction implies an inhibitory function of NS5A which might be released through stronger dimerization subsequently enhancing replication. In JFH1, both helicase and polymerase already had an intrinsically improved enzymatic activity allowing the virus to circumvent the ReED’s negative regulatory function. Thus, the discovery of a so far overlooked NS5A dimerization site helps to explain how HCV regulates its genome replication rate.

## Materials & Methods

### Cell culture

Clones of the immortalised hepatoma cell line Huh7-Lunet ectopically expressing SEC14L2 or SEC14L2 and T7 RNA polymerase were described previously (26). Using pWPI based plasmids, Huh7-Lunet SEC14L2 clones expressing GLT1 NS3-5A, GLT1 NS5A, Con1 NS5A, Con1 NS5A GLT1 ReED or just an empty puromycin cassette were generated for this study. They were cultured in Dulbecco’s modified Eagle’s medium (Gibco), supplemented with 1% (v/v) non-essential amino acids, 2 mM L-glutamine, 100 U/ml penicillin, 100 mg/ml streptomycin, 10% (v/v) heat-inactivated fetal-calf serum (FCS) and 5 µg/ml blasticidin, 1 mg/ml G418 (T7 overexpression) or 3 µg/ml puromycin (viral protein overexpression). All cell lines were regularly tested to check they were free of mycoplasma contamination using the MycoAlert Mycoplasma Detection kit (Lonza).

### Generation of stable cell lines

To generate stable cell lines, lentiviral particles were produced by transfecting HEK 293T cells with pWPI vectors using polyethyleneimine (PEI). The transfection complex was prepared by combining 6.42 μg of the pWPI vector, 6.42 μg of pCMV Δ8.91, and 2.16 μg of pMD2.G in 385 μl of Opti-MEM (Gibco). This solution was then added to a mixture of 45 μl PEI and 355 μl Opti-MEM. After vortexing and a 20 min incubation at room temperature, the mixture was applied to 5×10^6^ HEK 293T cells that had been seeded in 10 cm dishes 24 h prior. The culture medium was refreshed after 6 h. Following a 48 h incubation, the viral supernatant was collected, passed through a 0.45 μm filter, and transferred to the target cells (5×10^6^ HEK 293T or 6.4×10^5^ Huh7-Lunet SEC14L2 cells, seeded the previous day). After 24 h of transduction, the viral medium was replaced with fresh medium supplemented with 1.5 μg/ml puromycin to select for successfully transduced cells.

### Cloning

DNA fragments for cloning were generated via PCR using the PhusionFlash High-Fidelity Master Mix (Thermo Fisher Scientific) and/or via digest with restriction enzymes (New England Biolabs, Thermo Fisher Scientific). DNA fragments were combined either using the NEBuilder HiFi DNA assembly cloning kit (New England Biolabs) or the T4 DNA ligase (Thermo Fisher Scientific). Correct DNA sequence of the resulting plasmid was confirmed via Sanger or Oxford Nanopore sequencing (Microsynth AG). All plasmids used and generated during this study can be found in Table S1.

### In vitro transcription (IVT)

To prepare the templates, DNA plasmids were linearized using restriction enzymes targeting sites immediately downstream of the 3’UTR. The linearized DNA was then purified using the NucleoSpin Gel and PCR Clean-up kit (Macherey-Nagel). The IVT reaction was assembled by combining 10 μg of linearized DNA with 6 μl of T7 polymerase (homemade), 12.5 μl of rNTP solution (25 mM each), and 20 μl of 5x RRL buffer (1 M HEPES (pH = 7.5), 1 M MgCl_2_, 1 M Spermidine, and 1 M DTT). After adding 100 U of rRNasin RNase inhibitor (Promega), the volume was adjusted to 100 μl with nuclease-free water and the mixture was incubated overnight at 37 °C. The following day, template DNA was digested by adding 20 U of RQ1 DNase (Promega) for 1 h at 37 °C. For RNA purification, 420 μl dH_2_O, 60 μl 2 M sodium acetate (pH = 4.5) and 400 μl water-saturated phenol (pH < 5) were added. The mixture was vortexed, chilled on ice for 10 min, and centrifuged at 13,000×g (4 °C) for 10 min. The aqueous supernatant was recovered and mixed with one volume of chloroform, followed by centrifugation for 10 min at 13,000×g. The resulting supernatant was then mixed with 0.7 volumes of isopropanol and centrifuged again at 13,000×g for 10 min to precipitate the RNA. Finally, the pellet was washed with 70% ethanol, resuspended in dH_2_O, and the structural integrity of the RNA was verified via agarose gel electrophoresis.

### Electroporation

To transfect hepatoma cells with IVT RNA, the following electroporation procedure was employed. To prepare the cells, they were detached from the culture dish and washed once with PBS. They were then resuspended in Cytomix (120 mM KCl, 0.15 mM CaCl_2_, 10 mM K_2_HPO_4_/KH_2_PO_4_ (pH = 7.6), 2 mM EGTA, 25 mM HEPES, 5 mM MgCl_2_, plus freshly added 2 mM ATP and 5 mM glutathione) to a density of 10^7^ cells/ml. For electroporation, a 200 μl aliquot of this suspension (2×10^6^ cells) was mixed with 2.5 μg of the respective RNA in a 0.2 cm gap-width cuvette. Electroporation was performed using a BioRad GenePulser system set to 975 μF and 166 V. Post electroporation, cells were resuspended in 6 ml DMEM and seeded into 12-well plates. 4 h and 24 h timepoints: 1 ml of the cell suspension per well. 48 h and 72 h timepoints: 0.5 ml of suspension diluted with 0.5 ml of fresh medium per well. The cells were maintained at 37 °C in a 5% CO_2_ environment.

Drug treatments were performed 4 h after electroporation either the low dose NS5A inhibitor treatment with 1 μM Pibrentasvir (MedChem Express) or 10 pM Daclatasvir (Bristol-Myers Squibb) or varying concentrations of Alisporivir (Debiopharm).

### Luciferase assay

Samples for luciferase assay were harvested by washing with PBS and lysis in 200 µl lysis buffer (1% Triton X-100, 25 mM glycyl glycine, 15 mM MgSO_4_, 4 mM EGTA, and freshly added 1 mM DTT) per well. For firefly luciferase activity measurements, 80 µl lysate per technical replicate were mixed with 350 µl assay buffer (25 mM glycyl glycine, 15 mM K_3_PO_4_ (pH = 7.8), 0.15 M MgSO_4_, 4 mM EGTA (pH = 7.8) and freshly added 1 mM DTT, 2 mM ATP). During the subsequent luminescence measurement, a Lubat LB9510 tube luminometer (Berthold Technologies) added 100 µl of a luciferin solution (0.2 mM luciferin, 25 mM glycyl glycine) into the sample and then detected the signal for 20 s.

### NS5A dimerization assay

To quantify NS5A dimerization, for each variant two constructs either tagged with the small bit (sBit) or the large bit (lBit) which are two protein fragments that form nano luciferase when interacting were created. Huh7-Lunet T7 SEC14L2 cells were co-transfected with two plasmids, one expressing the sBit tagged and one expressing the lBit tagged protein. Transfection was performed by combining 0.5 μg of each plasmid with 100 μl of Opti-MEM (Gibco) and 3 μl of TransIT-LT1 (Mirus Bio), vortexing the mixture and incubating it for 30 min at RT. Then, 1.4 ml medium were added. For drug treatments, medium containing the respective concentration of Pibrentasvir (MedChem Express) was used. Per well of a 96-well plate, 150 μl of the resulting mixture was added to 1*10^4^ Huh7-Lunet T7 SEC14L2 cells seeded the day prior. After 24 h, cells were washed with PBS, lysed with 50 μl luciferase lysis buffer and stored at −20 °C. Luciferase assay was performed using the Nano-Glo luciferase assay system (Promega) with a Mithras LB 940 Plate Luminometer (Berthold Technologies).

### RdRp activity assay

Activity of recombinant Con1 NS5B or JFH1 NS5B GND (45) was determined by using the PicoGreen dye generating a fluorescent signal upon binding of dsRNA replication intermediates (44). The assay was performed in 96-well chimney black plates (Greiner) by combining 1 μg poly(C) RNA template, 0.5 mM rGTP (Jena Biosciences), 2.5 mM MnCl_2_, 5 mM NaCl, 5 mM DTT and 20 mM Tris-HCl (pH = 7.5) in a volume of 25 μl. The reaction was initiated by addition of 50 nM RdRp followed by incubation at 37 °C for 30 min and ended by addition of 175 μl PicoGreen (Invitrogen) diluted 1:350 in TE buffer. This mixture was incubated for 5 min at room temperature before measuring fluorescence intensity at 485 nm (excitation) and 520 nm (emission) with an Infinite M200 pro plate reader (Tecan).

### Immunoprecipitation (IP)

One day before transfection, 4×10^6^ Huh7-Lunet T7 SEC14L2 cells were seeded in a 175 cm² dish. The next day, a mixture of 15 μg of the respective plasmid DNA, 90 μl polyethyleneimine (PEI) (pH = 7.0) and 2 ml Opti-MEM (Gibco) was produced. It was incubated for at least 15 min at RT and then added to the cells. After 4 h, the medium was changed. 24 h after transfection, medium was removed, 1 ml PBS was added, cells were scratched off and transferred into a new tube. Next, they were centrifuged at 6000×g for 2 min, supernatant was removed and 2 ml lysis buffer (50 mM Tris/HCl (pH = 7.5), 5% glycerol, 0,5% 4-Nonylphenyl-polyethylene glycol (NP-40) (PanReac AppliChem), 1.5 mM MgCl_2_, 100 mM NaCl, EDTA-free Protease Inhibitor Cocktail (Roche)) were added. This was incubated on ice for 30 min and then centrifuged for at least 30 min at 13,000×g at 4 °C. 20 μl anti-HA magnetic beads (Thermo Scientific) were equilibrated twice with 1 ml lysis buffer. Afterwards, the supernatant of the cell lysis was added and incubated overnight at 4 °C with the beads while moving. The beads were washed three times with wash buffer (lysis buffer without NP-40 and protease inhibitor), dried and frozen.

### Mass spectrometry

For whole-cell proteomics (FP-MS), cell pellets (Huh7 cells) were lysed in guanidinium chloride buffer (6 M guanidinium chloride, 10 mM tris(2-carboxyethyl) phosphine (TCEP), 40 mM chloroacetamide (CAA), 100 mM Tris-HCl pH 8), boiled at 95 °C for 10 min and sonicated (4 °C, 15x 30 s on and 30 s off high setting, Bioruptor Plus, Diagenode). Protein concentration was determined by BCA assay (Pierce) according to the manufacturer’s instructions and adjusted to 50 µg. For affinity purification (AP-MS), proteins were denatured, reduced, and alkylated in 200 µl buffer of 0.6 M guanidinium chloride, 1mM TCEP, 4 mM CAA, 100 mM Tris-HCl pH 8. Protein digestion for FP-MS and AP-MS samples was performed by adding 0.5 µg LysC (WAKO Chemicals) and 0.5 µg trypsin (Promega) at 30 °C for 16 h. For peptide purification, we used StageTips made with three layers of C18 Empore filter discs (3M) and the eluted peptides were resuspended in a solution of 2% acetonitrile and 0.3% trifluoroacetic acid (TFA). All samples were measured on a Thermo LTQ Orbitrap XL Hybrid FT mass spectrometer (Thermo Fisher Scientific) coupled online to an EASY n-LC 1000 HPLC system (Thermo Fisher Scientific). The liquid chromatography setup consisted of a 20 cm reverse-phase analytical column (75 µm diameter, 60 °C; ReproSil-Pur C18-AQ 1.9 µm resin; Dr. Maisch) and the peptides were separated using a 120 min gradient at a flow rate of 300 nl/min, and a binary buffer system consisting of buffer A (0.1 % (v/v) formic acid in water), and buffer B (80 % (v/v) acetonitrile, 0.1 % (v/v) formic acid in water). The mass spectrometer was operated in data-dependent mode to automatically switch between full scan MS and MS/MS acquisition. The nanospray ion source was operated at 2.4 kV, heated capillary temperature set at 200 °C. Full scan MS1 spectra were recorded from 300 to 1,700 m/z and 60,000 resolution (at m/z 400). The precursor isolation window was set to 2.0 m/z and the normalized collision energy 35.0. For protein identification and quantitation, MS data files were processed with MaxQuant (version 2.0.1.0.) using the default settings and label-free quantification (LFQ) (LFQ min ratio count 2, normalization type classic) enabled. Spectra were searched against forward and reverse sequences of the reviewed human proteome (UP000005640_9606), hepatitis C virus genotype 1a (UP000000518), and supplemented by HA tag and single viral protein variants sequences for AP-MS. Values were log2-transformed and protein groups only identified by site, reverse matches and exclusion of contaminants from analysis. Results for all tested factors can be found in Table S2.

### AlphaFold modelling

All models were generated using AlphaFold 2.3 (41) with 12 recycles via a local implementation of ColabFold 1.5.5 (77). The AlphaFold-ptm model was used for monomeric predictions, whereas the AlphaFold-multimer-v3 (78) model was used for ReED dimeric modeling. Multiple sequence alignments were generated using the MMseqs2 server with the UniRef30 (79) and environmental databases (80). For each target, five models were generated. Amber relaxation was not performed.

### Molecular dynamics simulations

All-atom molecular dynamics simulations of the monomeric and dimeric AlphaFold models of a 33-residue ReED segment were prepared and simulated in Gromacs 2022.3 (81) on CPUs. The systems were minimized in vacuum over 20,000 steps using a steepest descent algorithm prior to assembly with the Amber 99SB-ILDN force field (82) and TIP3P water (83) in a cubic box with a minimum distance of 2 nm between the protein and the edge of the box. Charge neutrality was achieved by the addition of Na^+^ ions. The assembled systems were then minimized over 1000 steps, this time using a conjugate gradient algorithm, integrating a steepest descent every 50 steps. Afterwards, they were equilibrated to a temperature of 300 K for 500 ps under NVT conditions (constant volume and temperature) employing the Berendsen thermostat (84) with a time constant of 0.1 ps and applying positional restraints on the peptide. Next, the systems underwent pressure equilibration to 1.0 bar for 500 ps under NPT conditions (constant pressure and temperature) governed by an isotropic Berendsen barostat with a 1.0 ps time constant and a compressibility factor of 4.5×10^−5^ bar^−1^, keeping the positional restraints on the peptide. Restraints were removed for production, and the systems were simulated over 200 ns. Both equilibration and production runs were carried out with an integration timestep of 2 fs. The V-rescale thermostat (85) and an isotropic Parrinello-Rahman barostat (86) were applied during production, keeping the parameters from equilibration, except for an updated barostat time constant of 2.0 ps. The LINCS algorithm (87) was employed to constrain bonds between heavy atoms and hydrogens. The Verlet cut-off scheme (88) was adopted for managing long range interactions, with a threshold distance of 1 nm. Secondary structure was computed with MDAnalysis (89) and the simulation snapshots were rendered on PyMOL (90).

### Purification of recombinant ReEDs

The DNA sequences coding for Con1, GLT1 and FCH3 LCS1D2 were synthesized by Genecust with codon optimization for expression in *E. coli*. These sequences were then inserted into the pET24b vector between the *Nco*I and *Xho*I restriction sites. The recombinant proteins contained a C-terminal 6xHis-tag. Each plasmid was introduced into *E. coli* BL21(DE3) cells, which were then grown in an LB medium supplemented with kanamycin. Following induction with 0.4 mM IPTG, protein expression was performed at 37 °C for 4 h. The cells were then harvested by centrifugation and resuspended in 40 ml of lysis buffer (50 mM phosphate buffer, pH 7.8; 300 mM NaCl; 0.5 mM DTT; protease inhibitor cocktail). This was supplemented with both DNase I (150 µl at 6 mg/ml) and RNaseA (50 µl at 40 mg/ml), after which the cells were lysed by sonication. After centrifugation at 20,000×g for 30 min at 4 °C, the supernatant was loaded onto a 1 ml HisTrap HP column (Cytiva) that had been equilibrated with Buffer A [50 mM sodium phosphate buffer at pH 7.8 and 300 mM NaCl]. After extensive wash with Buffer A, the recombinant LCS1D2 proteins were eluted using a 0-100% gradient of Buffer B (Buffer A supplemented 300 mM imidazole) over 10 column volumes. The fractions containing LCS1D2 were selected, based on SDS-PAGE analysis, pooled, and the buffer was exchanged for 50 mM sodium phosphate buffer pH 7.8 with 150 mM NaCl. The proteins were concentrated using a centrifugal concentrator (Vivaspin Turbo, Sartorius), then flash-frozen in liquid nitrogen and stored at −80 °C until required.

### Size exclusion chromatography

Recombinant LCS1D2 proteins were analyzed by size-exclusion chromatography using a Superdex 75 Increase 10/300 GL column (Cytiva), which was equilibrated in 20 mM Tris.Cl pH 8.0, 200 mM NaCl. 100 µl LCS1D2 samples (at ∼50 µM) were injected onto the column using a 100 µl loop with a flow rate of 0.5 ml/min.

## Acknowledgements

We thank R. Klein and U. Herian for excellent technical assistance. For the generous gift of plasmids and antibodies, we are grateful to J. Bukh (J6), T. Wakita (JFH1) and C. Rice (α-NS5A 9E10). We thank J. Becker and V. Gonçalves Magalhães for helpful discussions.

## Notes

### Competing Interest Statement

The authors have declared no competing interest.

